# Haploinsufficient maternal effect of Epidermal Growth Factor Receptor A mutation in zebrafish

**DOI:** 10.1101/745018

**Authors:** Margherita Ciano, Paul R. Kemp, S. Amanda Sathyapala, Simon M. Hughes

## Abstract

Generation of viable offspring depends both on genetic and environmental factors of both mother and child. Analysis of a likely amorphic allele of the zebrafish *epidermal growth factor receptor a* (*egfra*) gene revealed that heterozygous females were infertile due to death of all fertilized eggs during embryonic and early larval life with cardiac, tail and other defects. Comparison of the severe dominant maternal effect with previous studies using pharmacological inhibitors of Egfrs or antisense morpholino injection indicate that a normal level of maternal Egfra is required for viability of offspring both during egg development and in the embryo after fertilisation. As heterozygous mothers were not fertile, the homozygous zygotic *egfra^kg134^* phenotype could not be analysed. Heterozygous *egfra^+/kg134^* males crossed to wild type females produced fully viable offspring, among which *egfra^+/kg134^* individuals had increased slow muscle but no functional motility defect. Our findings suggest that Egfra activity is crucial for early development both before and after fertilisation and are likely to constitute a rare example of a haploinsufficient maternal effect in a species lacking imprinting.

## Introduction

Regulation of fertility is important for humans and food production. Development of oocytes and early embryos depends on the contribution of protein and RNA encoded by the maternal genome prior to the final meiotic division to yield the mature gamete. Maternal effect genetic screens have been a powerful tool in unravelling early embryo development across vertebrates (Marlow, 2010). A particularly important aspect of oocyte development is its interaction with other cells within the ovary, such as follicular or nurse cells, which are mediated by a wide range of reciprocal extracellular signals (Richani and Gilchrist, 2018). Mutation of genes required for such signals have been shown to have effects on the developing oocyte or embryo when the mother is homozygous for the mutation, irrespective of ultimate embryonic genotype (Marlow, 2010). In general, however, maternal effect mutations are recessive, such that heterozygous females yield offspring that develop normally unless the offspring inherit an allelic combination causing a zygotic phenotype. Here we describe dominant maternal effect caused by mutation in the zebrafish *egfra* gene.

Epidermal growth factor (EGF) signalling through EGFR receptors of the ErbB family is of central importance in human reproduction (Richani and Gilchrist, 2018). In mammals, several EGFR ligands and the EGFR itself are required for oocyte maturation and ovulation (Hsieh et al., 2007). The EGFR is not thought to be present on the oocyte itself, but acts via the surrounding cumulus cells, in which EGFR signalling triggers expansion of the cumulus and cumulus-derived extracellular matrix and promotes oocyte maturation through downregulation of cyclic nucleotides that are shared with the oocyte through gap junctions and suppress meiotic resumption. EGFR signaling within the follicle also enhances oocyte ATP production and subsequent oocyte developmental competence (that is, the ability of an oocyte to yield, upon fertilization, an embryo that successfully develops to term; (Richani and Gilchrist, 2018) and references therein). Indeed, levels of the EGFR ligand Amphiregulin in follicles correlate with infertility and successful IVF treatment (Ambekar et al., 2015; Huang et al., 2015). Strikingly, in mouse, maturation of EGFR signalling competence in cumulus cells is itself triggered by oocyte-derived signals, supporting the view that stepwise maturation of the follicle involves continual crosstalk between oocyte and its support cells that modulate EGFR signaling (Richani and Gilchrist, 2018). It is, therefore, of great importance to understand the roles of EGFR in oocyte maturation and acquisition of developmental competence.

EGF signalling has been extensively characterized in cultured cells and in mammals, in large part due to its importance in a variety of cancers (https://omim.org/entry/131550). During murine development, *Egfr* is an essential gene; null mutation leads to genetic background-dependent prenatal death due to placental insufficiency, and additional lung, epidermis, whisker and eye defects (Miettinen et al., 1995; Sibilia and Wagner, 1995; Threadgill et al., 1995). Numerous other mutant alleles affect hair and skin and cause defects in a variety of organs including heart and lung. However, genetic loss of function analysis in vertebrates beyond the mouse has not been performed.

In the zebrafish, *egfra* function has been implicated in cardiac outflow tract formation, aorta thickness, adult intestinal adaptation and ovarian follicle development (Aizen and Thomas, 2015; Goishi et al., 2003; Peyton and Thomas, 2011; Schall et al., 2015; Wang and Ge, 2004; Zhao and Lin, 2013). In embryos/larvae, inhibition of zebrafish Egfra with either AG1478 or PK166, two drugs that inhibit mammalian EGFRs, leads to death from pericardial oedema, an effect that can be phenocopied by *egfra* antisense morpholino injection (Goishi et al., 2003). We previously described generation of the putative null *egfra^kg134^* allele in the zebrafish in which a frameshift 12 amino acids from the N-terminus of the protein leads to a severe truncation. Heterozygous *egfra^+/kg134^* larvae have a mild increase in slow muscle but are viable, whereas homozygous mutants die early in larval life (Ciano et al., 2019). However, in our initial report, more extensive analysis of the genetics of the *egfra^kg134^* allele was not reported.

Here we analyse the genetics of the *egfra* loss of function mutation in more detail and report the striking finding that the *egfra^kg134^* allele shows a maternal effect in heterozygous mothers, such that no offspring survive beyond early larval life, irrespective of zygotic genotype. In contrast, heterozygous males yield viable heterozygous offspring of both sexes.

## Results

The *egfra* mutant allele *egfra^kg134^* generated by CRISPR/Cas9 genome editing has a small indel that leads to a frameshift after 12 amino acids in the first coding exon. This results in termination of the resultant polypeptide with a 70 amino acid nonsense tail at a normally out-of-frame stop codon in the second coding exon (Ciano et al., 2019). The predicted protein lacks essential EGF binding regions, the transmembrane domain and the intracellular tyrosine kinase of EGFR and is therefore highly unlikely to have any residual positive or dominant negative function. Further evidence for lack of a dominant negative effect is a) that the mutant allele causes nonsense-mediated mRNA decay, so little truncated protein will be produced (Ciano et al., 2019) and b) that heterozygous *egfra^+/kg134^* larvae do not show any of the severe defects reported in *Egfr* null mutant mice (Miettinen et al., 1995; Sibilia and Wagner, 1995; Threadgill et al., 1995).

### Fertilized eggs from *egfra^+/kg134^* heterozygous females die early

Heterozygous F1 *egfra^+/kg134^* fish were in-crossed to generate embryos homozygous for the mutation. However, all embryos died in the lays. Four F1 *egfra^+/kg134^* in-crosses between the single available adult F1 carrier *egfra^+/kg134^* female (designated α) and different *egfra^+/kg134^* F1 males were performed. No embryos survived beyond 4 days post fertilisation (dpf). Most deaths occurred between 1 and 2 dpf (Fig. 1A). Genotyping of resultant F2 embryos failed to reveal differences based on zygotic genotype.

**Figure 1.**
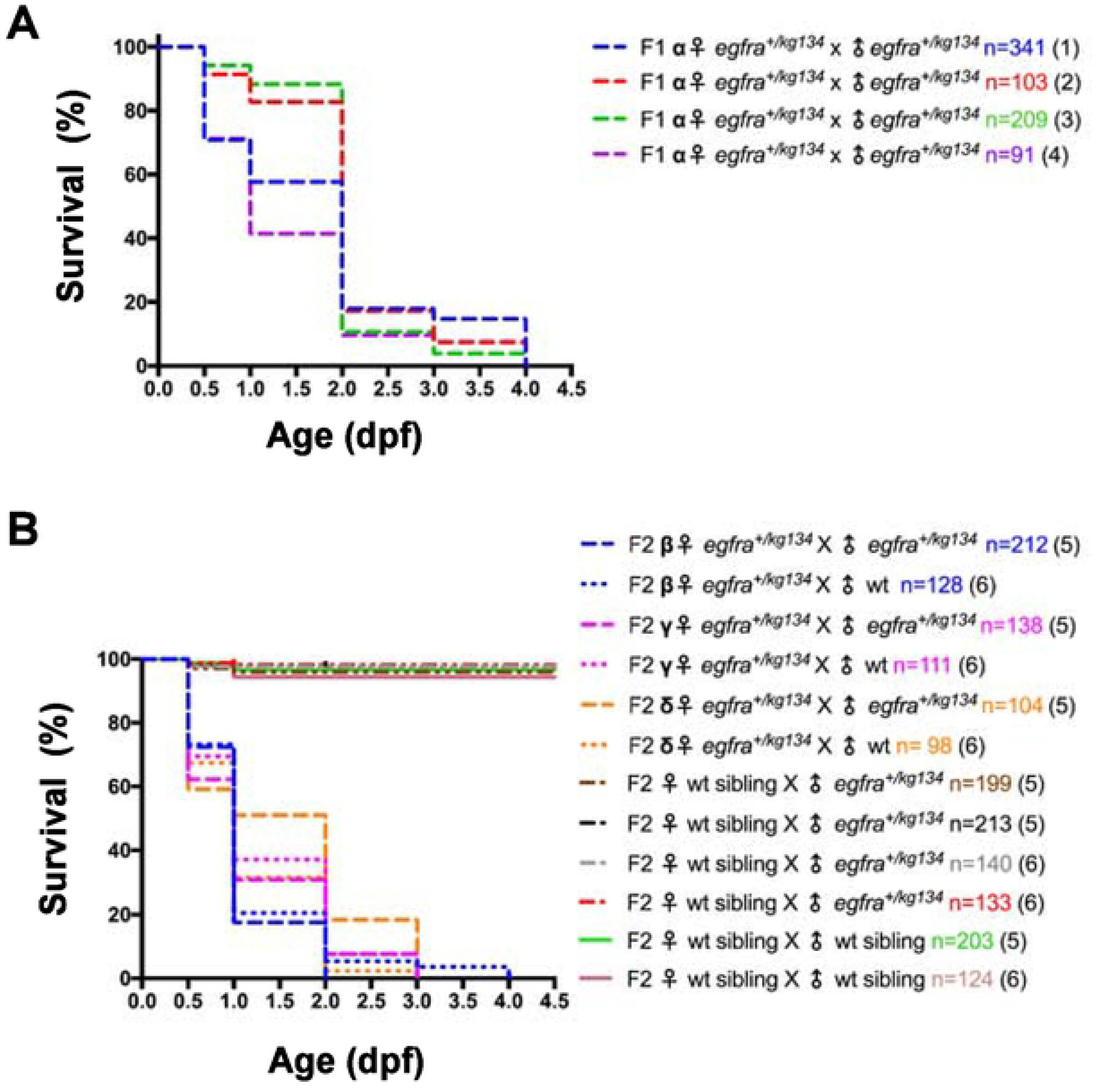
Survival of embryos from *egfra^+/kg134^* and sibling females. **A, B.** Survival curves for embryos from the indicated crosses. The single F1 female obtained (designated α) was crossed repeatedly and survival of progeny monitored every twelve hours (A). Three separate F2 *egfra^+/kg134^* females (designated β, γ and δ) were also tested (B). Note that lays from wild type (wt) sibling females crossed with wt or heterozygous *egfra^+/kg134^* males survive, whereas lays from heterozygous *egfra^+/kg134^* females die, irrespective of the partner male genotype.

F2 generations were bred by out-crossing carrier *egfra^+/kg134^* males to wild type females. After genotyping to identify *egfra^+/kg134^* F2 males and females, three females (designated β, γ and δ) and various males were obtained. Repeatedly, embryos laid by F2 *egfra^+/kg134^* females all died (Fig. 1B). In contrast, embryos laid from wild type *egfra*^+/+^ sibling F2 females developed normally and survived (Fig. 1B). Paternal genotype had no discernible effect; fertilisation of wild type-derived eggs by sperm from either wild type or *egfra^+/kg134^* F2 sibling males yielded viable embryos (Fig. 1B). An F3 generation was bred by out-crossing an *egfra^+/kg134^* F2 male to a wild type female. Female F3 *egfra^+/kg134^* carriers again failed to yield viable offspring (data not shown). The number of eggs produced by *egfra^+/kg134^* females was not significantly different from that produced by their wild type siblings. These data suggest that female zebrafish possessing only a single functional *egfra* allele are not fertile because they produce eggs that cannot develop normally.

### Heterozygous maternal effect causes abnormal embryo development

Maternal effect is often caused by the requirement of gene function in the egg or fertilized embryo from mRNA placed in the egg during oogenesis (Marlow, 2010). We performed in situ mRNA hybridization on newly-laid eggs and embryos prior to activation of the zygotic genome and observed *egfra* mRNA accumulation in the early embryos (Fig. 2A). This finding indicates that *egfra* mRNA derived from maternal chromosomes is accumulated in the egg.

**Figure 2.**
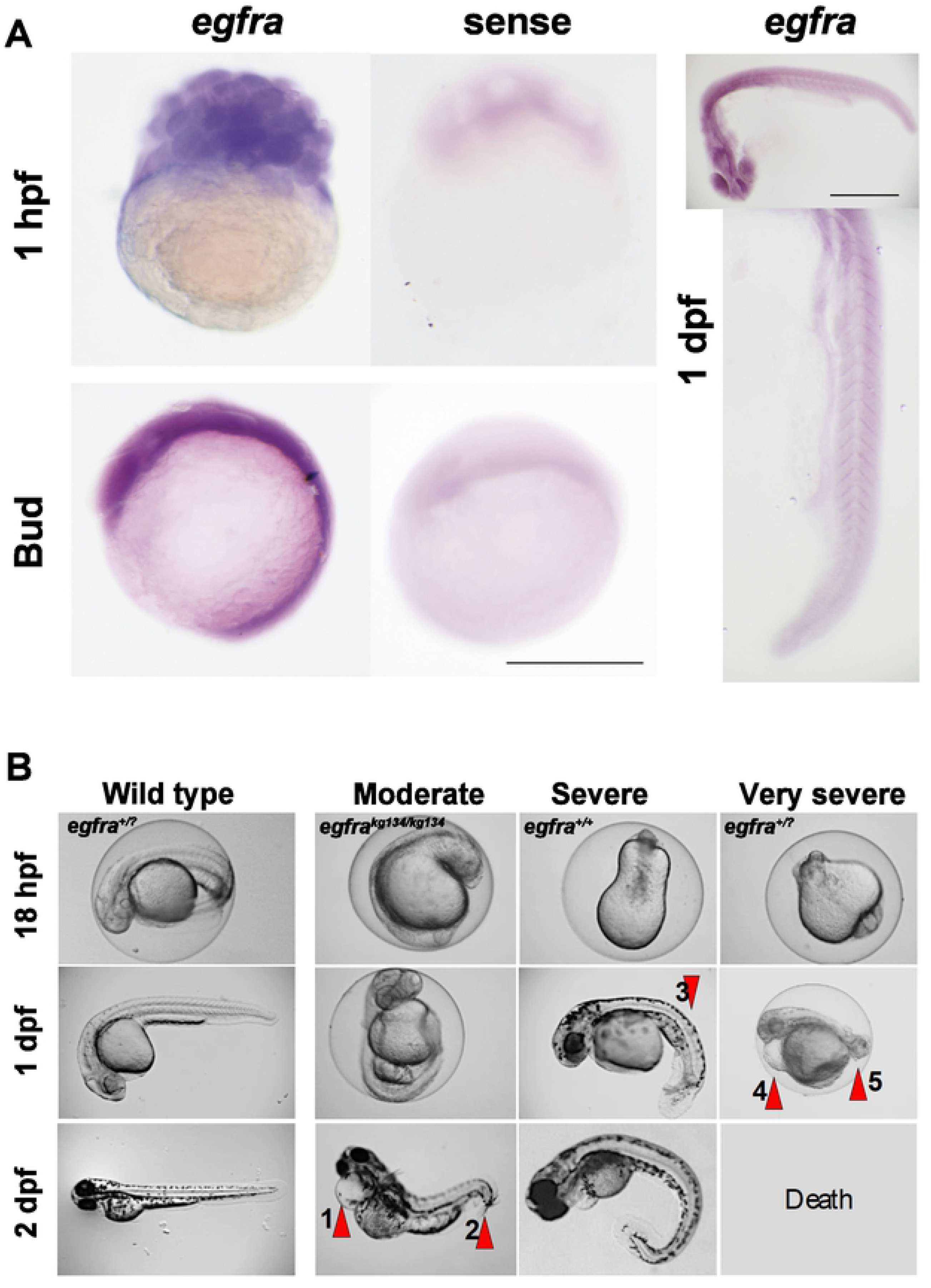
Heterozygous maternal effect may arise from maternal *egfra* expression in early embryos. **A.** In situ mRNA hybridisation for antisense (left) and sense (right) *egfra* probes at 32 cell stage, after gastrulation and at 1 dpf. Bars = 500 μm. Note abundant expression at 32 cells, prior to zygotic genome activation and widespread but declining mRNA until 24 hpf. **B.** Representative timelapse imaging of single progeny of a wild type sibling female (left) and three maternal effect phenotypes (right) from *egfra^+/kg134^* females crossed to *egfra^+/kg134^* males. As all embryos from the wild type female appeared wild type the progeny were not individually genotyped and are therefore designated *egfra^+/?^*. All embryos from the carrier female showed growth retardation, with a moderate group characterised by cardiac oedema (arrow 1) and curvature at the tail tip (arrow 2), of which a zygotic mutant is shown. A severe group presented with a beanshaped yolk at 18 hpf that, by the end of day 1, showed curvature of the trunk and tail (arrow 3), which persisted even after a day outside of the chorion; a zygotic wild type is shown. A very severe group had poor tailbud formation at 18 hpf, cardiac oedema (arrow 4) and failure of tail development (arrow 5) during day 1. Such embryos die by 2 dpf and the one shown was therefore not genotyped. All three phenotypes were also obtained from *egfra^+/kg134^* females crossed to *egfra^+/+^* males.

Fertilisation and egg activation rates appeared normal in clutches derived from *egfra^+/kg134^* females (Fig. 1). Chorion elevation was indistinguishable from wild type (data not shown). Although small numbers of unfertilized embryos were observed in some lays, as reflected by the slight drop in survival during the first dpf in some control lays (Fig. 1B), such unfertilized embryos were readily distinguished from fertilized embryos undergoing cleavage in lays from *egfra^+/kg134^* females. No defects in cleavage stage development were noted. Moreover, epiboly and gastrulation were not grossly defective (data not shown). By 18 hpf, however, almost all embryos in clutches from *egfra^+/kg134^* females were defective (Fig. 2B). Subsequently, a range of defects of variable severity appeared. Moderately affected embryos had a curved tail tip sometimes accompanied by cardiac oedema. ‘Severe’ embryos had unusual yolk shapes and showed altered tailbud morphology and poor yolk extension. Very severely affected embryos lacked a tail and showed signs compatible with widespread apoptosis (Fig. 2B). The severity of defects did not correlate with zygotic *egfra* genotype. For example, the moderately affected embryo shown in Fig. 2B was a zygotic mutant derived from an *egfra^+/kg134^* female crossed to an *egfra^+/kg134^* male. In contrast, more severely defective embryos could be *egfra^+/kg134^* or wild type, whether derived from a heterozygote in-cross or an *egfra^+/kg134^* female crossed to a wild type male (Fig. 2B and data not shown). We conclude that reduction of Egfra function derived from the mother causes a severe developmental defect.

### Behaviour of mutant and wild type larvae

To examine the role of zygotic *<u>egfra</u>* further, bearing in mind the increase in slow muscle fibres previously observed (Ciano et al., 2019), differences in burst swimming in response to touch were investigated in F3 *egfra^+/kg134^* and wt siblings. No significant difference in motility were observed (Fig. 3). Heterozygous *egfra^+/kg134^* and wt sibling larvae grew to adulthood equally well and were indistinguishable as adults (data not shown).

**Figure 3.**
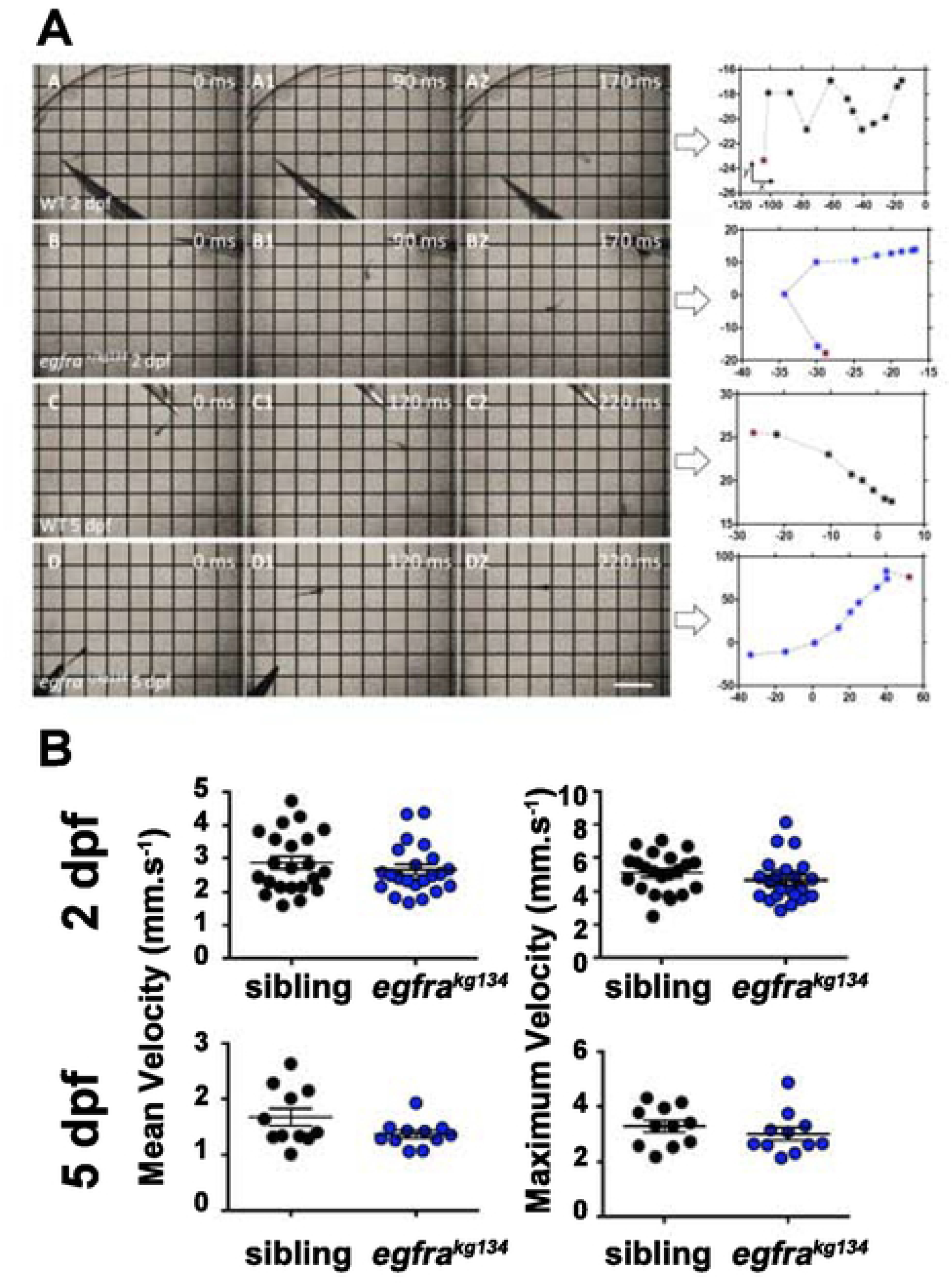
F3 *egfra^+/kg134^* and wild type siblings swim similarly. **A.** Images extracted from burst response videos of F3 2 dpf sibling wt (A-A2) and *egfra^+/kg134^* (B-B2) larvae and 5 dpf sibling wt (C-C2) and *egfra^+/kg134^* (D-D2) larvae. Timepoints are shown in milliseconds (ms). Arrows indicate the plot of the tracking analysis with the *X* and *Y* coordinates, the red asterisk in the plot indicates the start point of the fish, black symbol: WT fish, blue symbols: *egfra^+/kg134^* fish. Bar 5 mm. **B.** Diagram to illustrate how measurement of Velocity was performed. **C.** Motility of 2 dpf and 5 dpf F3 *egfra^+/kg134^* and wt sibling larvae were compared. No difference was found at either age (2 dpf N=22/group from 2 crosses, mean velocity p=0.68 and maximum velocity p=0.12; 5 dpf N=11/group from 2 crosses, mean velocity p=0.21 and maximum velocity p=0.32, Mann-Whitney U test).

## Discussion

The data presented strongly suggest that *egfra* has an essential role on early embryonic development and that the level of the expressed Egfra receptor is crucial. As we have only isolated and characterised a single mutant allele, we cannot rule out the possibility that the sgRNA used generated an off-target mutation closely-linked to the *egfra^kg134^* allele on chromosome 2. We think this unlikely because i) no close matches with adjacent PAM sites are present nearby on chromosome 2 (CRISPRdirect shows zero 20 bp or 12 bp matches and the nearest 7/8 bp seed match with an adjacent PAM site are 3 Mb distal and 1.5 Mb proximal on chromosome 2 and neither lies in or near a known coding region), ii) two other lines with F1s carrying distinct *egfra* mutant alleles generated with the same sgRNA did not yield fertile F1 females and were discarded before the nature of the maternal effect became apparent, iii) *egfra* is expressed in the developing oocyte, egg and early embryo, consistent with a maternal effect (Goishi et al., 2003), and iv) Egfr has previously been implicated in oocyte development in zebrafish (Peyton and Thomas, 2011; Van Der Kraak and Lister, 2011).

The correct level of the Egfra receptor appears to be important for zebrafish fertility. We previously reported that *egfra* mRNA is reduced by 50% in *egfra^+/kg134^* fish (Ciano et al., 2019), consistent with nonsense mediated decay of the mutant mRNA, bearing in mind that the minor sequence change in exon 1 is not predicted to affect transcription, splicing, or translational initiation. It is therefore likely that during some or all of egg development and early embryonic life, Egfra protein level in the developing gamete/embryo may be reduced. It has been suggested that oocyte maturation depends on the balance of Egfr (also known as Erbb1) and other Erbb receptors, such as Erbb2, that can heterodimerise (Aizen et al., 2018). Other work has suggested a role for Egfr activity in the follicle cells during oocyte maturation in zebrafish (Wang and Ge, 2004) and ovulation in mice (Ashkenazi et al., 2005; Hsieh et al., 2007; Jamnongjit et al., 2005). Treatment of female fish with the environmental dioxin pollutant TCDD reduces both fertility and *egfra* level in the ovary (Heiden et al., 2008). However, we observe normal egg abundance, mating behaviour, fertilisation rates and oocyte activation from *egfra^+/kg134^* females. Moreover, early stages of cleavage and epiboly also appear normal. Our *egfra^kg134^* maternal effect mutation has late and pleiotropic effects. We therefore hypothesise that the relevant period for the requirement for the correct level of Egfra may extend into early embryonic life. But an alternative possibility is that lack of appropriate EGFR signalling during oocyte development reduces oocyte quality such that subsequent embryo development is compromised. The fact that injection of *egfra-* targeting morpholino antisense oligonucleotides into 1-cell stage embryos does not phenocopy the *egfra^+/kg134^* maternal effect (Goishi et al., 2003), suggests the maternal Egfra protein functions before fertilisation and/or may be stockpiled in the egg.

Our data add an evolutionary perspective to the increasing understanding of the importance of EGFR signalling in mammalian oocyte development (Richani and Gilchrist, 2018). Our data show that reduction of maternal Egfra function generates poor quality eggs, irrespective of zygotic genotype. Despite fertilisation and apparently normal cleavage, embryos from *egfra^+/kg134^* mothers develop poorly and die. In humans and other mammals, oocyte maturation and high oocyte developmental competence depend on EGFR signalling within the follicle that acts on cumulus cells and may affect in vitro maturation procedures that could offer significant benefit to women with polycystic ovary syndrome or cancer (Richani and Gilchrist, 2018). Such signalling is thought to enhance metabolic capacity and regulate post-transcriptional events that prepare oocytes for their future developmental role in the embryo (Chen et al., 2013; Richani et al., 2014; Sugimura et al., 2015). Our finding that EGFR signalling controls developmental competence in zebrafish suggests that EGFR signalling is an ancient and important pathway in oogenesis that is conserved across vertebrate evolution.

Treatment of zebrafish embryos with Egfr-blocking drugs causes a variety of defects (Budi et al., 2008; Goishi et al., 2003), but those described are not as severe as the early death we observe. This supports the view that Egfra activity is required both prior to egg laying for correct egg development and after egg laying for later developmental steps. Morpholino knockdown of Egfra leads to cardiac and vascular abnormalities that have some similarities to the variable penetrance cardiac oedema we observe (Goishi et al., 2003). Altered Egfra activity has also been described to cause biliary atresia and to alter gut repair (Ningappa et al., 2015; Schall et al., 2015), processes that may contribute the phenotypes we observe, but are unlikely to cause the gross embryonic defects and early death.

We also report further characterisation of the phenotype of *egfra^+/kg134^* fish, which have extra slow muscle fibres in their somitic myotome (Ciano et al., 2019). These *egfra^+/kg134^* fish did not change speed of swimming triggered by a touch, a validated physiological measure that is fast fibre-dependent (Naganawa and Hirata, 2011). As the level of Hedgehog (Hh) signaling also controls the number of slow muscle fibres in zebrafish somite, it is possible that the EGF-repeat containing protein Scube2, which is required for normal slow fibre formation could provide a link to EGF/Egfr function (Blagden et al., 1997; Hollway et al., 2006). However, because Scube2 lacking the EGF domains can rescue Hh palmitoylation and signaling in vitro and in vivo (Creanga et al., 2012), it is unclear if reduction of Egfra function affects slow myogenesis through modulating Hh signalling. Alternatively, Egfra function may suppress the later Hh-independent additional slow myogenesis, as patterned expression is observed in the somite after 1 dpf (Fig. 2B)(Barresi et al., 2001; Ciano et al., 2019). To date we have been unable to discern a zygotic functional phenotype in *egfra^+/kg134^* fish. Unfortunately, *egfra^kg134^* homozygote mutants do not survive well or long enough for meaningful functional studies.

Finally, the most notable aspect of our findings is the existence of such a highly penetrant heterozygous maternal effect that we attribute to haploinsufficiency. Female mice lacking one allele of *Egfr* do not show a similar infertility (Miettinen et al., 1995; Sibilia and Wagner, 1995; Threadgill et al., 1995). To our knowledge, few dominant maternal effect genes have been described, although many may have been discarded in the extensive screens for recessive zygotic and maternal effect mutations (Marlow, 2010). Rare known examples of maternal haploinsufficiency are the unmapped *mel-23^ct45^* allele in *C. elegans* and the Minute allele *RpS17^4^* in *D. melanogaster* (Boring et al., 1989; Mains et al., 1990). In mammals, genetics mimicking maternal haploinsufficiency can be observed after uneven X-chromosome inactivation or in imprinted genes where the paternal allele is not expressed. However, zebrafish have neither sex chromosomes, nor parental imprinting. Moreover, because we used genome editing to create and select *egfra^kg134^* as a likely loss of function *egfra* allele, we suggest that the apparent dominant maternal effect is caused by maternal haploinsufficiency of Egfra. Although to date searches of OMIM and extant QTL data have failed to reveal linkage of human *EGFR* variation to infertility, a possible genetic influence on EGFR signalling in fertility in humans and other vertebrates should be borne in mind.

## Methods

### Zebrafish lines and maintenance

Zebrafish were reared at King’s College London on a 14/10 hours light/dark cycle at 28.5 °C with adults kept at 26.5°C, with staging and husbandry as described (Westerfield, 2000). Founder and subsequent generations were out-crossed to AB fish. Fish were genotyped by High Resolution Melt Analysis and/or DNA sequencing in the mutant alleles, following PCR amplification using primers indicated (Supplementary Table 1). Individual genotyped carrier *egfra^kg134/^+* females were kept in separate tanks to ensure reproducibility. All experiments were performed in accordance with licences held under the UK Animals (Scientific Procedures) Act 1986 and later modifications and conforming to all relevant guidelines and regulations.

### Imaging and in situ mRNA hybridization

ISH was performed as described (Ganassi et al., 2014). Briefly, fish were fixed in 4% paraformaldehyde (PFA) in phosphate-buffered saline (PBS) for 30 min or 3 hours at room temperature, stored in 100% methanol at −20°C and rehydrated in PBS prior to ISH. Digoxigenin-labelled probes were against *egfra* as described (Ciano et al., 2019) and embryos were imaged under a Leica MZ16F with Olympus camera.

### Motility Assay

Embryos touch response was measured on F3 larvae at 2 and 5 dpf and recorded with a Leica DFC490 colour camera at 500 frame per seconds, as previously described (Naganawa and Hirata, 2011). Each embryo was imaged in a petri dish of fish water under a Leica MZ16F microscope, the tail touched with forceps to trigger movement and behaviour recorded for 10s. Videos were analysed blind with Tracker software 4.97 (https://physlets.org/tracker/) and mean and maximum velocity calculated. DNA was subsequently extracted from each embryo for genotyping.

## Author contributions

SMH and SAS conceived the project and obtained finance. MC performed all experiments and analysis. PRK provided advice. SMH and MC wrote the paper with input from other authors.

## Acknowledgements

We are grateful to Giorgia Bergamin and all members of the Hughes lab for advice, and to Bruno Correia da Silva and his staff for care of the fish.

---

**Supplementary Table 1 Primers for genotyping**

HRMA primers 5’-CCCGATAGCTTACAAACGCA-3’ 5’-GCCGTTTCACAATAGTCCTACC-3’ Sequencing primers 5’-GGAGGAGGAGCTGTCAAAGT-3’ 5’-GCGATGTTCCCAAATCATTTTCC-3’

